# Rodents as Potential Reservoirs for Toroviruses

**DOI:** 10.1101/2025.06.12.659308

**Authors:** Jaime Buigues, Adrià Viñals, Raquel Martínez-Recio, Juan S. Monrós, Rafael Sanjuán, José M. Cuevas

## Abstract

Emerging zoonotic viruses pose a significant threat to global health. The order Nidovirales includes diverse viruses, such as coronaviruses, which are well known for their zoonotic potential. Toroviruses are a less-studied genus within Nidovirales primarily associated with gastrointestinal diseases in ungulates, although some evidence suggests their presence in humans. In this study, we report the discovery of a novel torovirus from a fecal sample of a dormouse (*Eliomys quercinus*) in Spain, which we named Dormouse torovirus (DToV). This represents the first complete genome of a rodent-associated torovirus. The 28,555-nucleotide genome encodes the six characteristic torovirus open reading frames, but these exhibit low amino acid sequence identity (44.3–86.3%) compared to other toroviruses, indicating that DToV likely represents a new viral species. Bayesian analysis of the ORF1b suggests that DToV diverged from known toroviruses approximately 1300 years ago. Moreover, the basal phylogenetic position of DToV suggests that rodents may represent a reservoir for this viral genus. Our findings expand the known torovirus host range, underscore their potential for cross-species transmission, and highlight the importance of continued surveillance of wildlife viruses.

**Importance:** Currently, there is concern about the potential emergence of new viruses from natural reservoirs. This has led to an increasing search taking advantage of new massive sequencing methodologies. In our study, we have identified a complete genome of a torovirus in a rodent species in Spain. Toroviruses are a group related to coronaviruses and cause gastrointestinal diseases in ungulates, such as cows or pigs. However, they have not been studied in depth, as there is no clear evidence of their ability to infect humans. Our results suggest that rodents may be the natural reservoir of toroviruses and underline their potential for cross-species transmission. Based on the results obtained from monitoring wildlife for viruses with zoonotic potential, the next step will be to implement new approaches in the laboratory to characterise the ability of these viruses to infect humans.

## 1. Introduction

Viral zoonoses have been responsible for wide-ranging human health impacts (Watsa 2020; Vora *et al*. 2023). This has led to the strengthening of pathogen surveillance programs to collect information on circulating viruses in human and wildlife populations (Watsa 2020). To this end, viral metagenomics has become the most widely used tool to characterize circulating viruses in different ecosystems (Cui *et al*. 2023). The search for wildlife viruses has focused on tropical areas, since land-use alterations, high wildlife diversity, and bush meat consumption, among other factors, are believed to increase disease emergence risk in these regions (Allen *et al*. 2017; Huong *et al*. 2020). However, it is important to extend this search to other regions, as some emerging viral diseases did not originate in tropical areas (Gibbs, Armstrong and Downie 2009; Han, Yu and Yu 2016). In Europe, for instance, animal-to-human viral transmission events include those of tick-borne encephalitis virus, Dobrava virus, and Granada virus, among others (Kallio-Kokko *et al*. 2005).

Wildlife reservoirs are responsible for most emerging infectious diseases (Cui *et al*. 2023), but certain animal taxa give rise to more zoonoses than others. Although bats seem to harbor a significantly higher proportion of zoonotic viruses than all other mammalian orders (Olival *et al*. 2017), rodents are the most speciose group, and some species live in close contact with humans, increasing zoonotic risk (Luis *et al*. 2015; Buigues *et al*. 2024). Dwellings and agricultural settings are among the highest-risk interfaces for zoonotic viral transmission, particularly from rodents (Kreuder Johnson *et al*. 2015). For instance, the house mouse has facilitated the worldwide transmission of viruses to sympatric species (Luis *et al*. 2015). In addition, rodent hantaviruses and mammarenaviruses are probably the original reservoir of embecoviruses, which gave rise to human coronaviruses OC43 and HKU1 (Cui *et al*. 2023).

RNA viruses, particularly those with envelopes, are more prone to cross-species transmission and zoonosis than other viruses (Geoghegan and Holmes 2018; Valero-Rello and Sanjuán 2022). A salient example of this group of viruses is the *Coronaviridae* family (order *Nidovirales*), which includes viruses that have repeatedly demonstrated their ability to switch hosts (Kane, Wong and Gao 2023). Much less known is the genus *Torovirus*, historically within the family *Coronaviridae*, but currently assigned as a unique genus to the subfamily *Torovirinae*, family *Tobaniviridae* (Walker *et al*. 2019), which includes three species: bovine torovirus (BToV), equine torovirus (EToV), and porcine torovirus (PToV), as defined by the International Committee on Taxonomy of Viruses (ICTV; https://talk.ictvo nline.org/taxonomy). Complete torovirus genome sequences have been found in several ungulates and a marsupial carnivore, whereas only partial sequences have been detected in humans (Ujike and Taguchi 2021), and there is no strong evidence for the existence of human toroviruses. Indeed, while toroviruses cause gastrointestinal disease in ungulates, no pathological associations have been observed in humans so far (Ujike and Taguchi 2021).

However, the current lack of knowledge about toroviruses makes it difficult to rule out their potential involvement in future zoonotic events (Ahmad *et al*. 2024). In this context, given the threat that some members of the Nidovirales group pose to human health, our study focused on identifying viruses classified in this order. To do so, a metagenomic analysis of fecal samples from small terrestrial mammals in Spain was performed. A novel species of torovirus, which we called Dormouse torovirus (DToV), was identified in a rodent species (*Elyomis quercinus*).

## 2. Material and methods

### 2.1. Study area and sample collection

Box-like traps (Sherman, Ugglan, and Mesh traps) were used to capture small terrestrial mammals from different locations in eight Spanish provinces (Alicante, Cantabria, Huelva, León, Madrid, Salamanca, Valencia, and Zamora; **Supplementary Table S1**). The sampling period spanned from March to November 2022. A total of 160 individuals were captured, for which species, sex, and age were determined before release, except in a few cases, where samples were taken, without capture, from burrows of previously identified species. Most of the individuals belonged to the order *Rodentia*, and a small number of cases belonged to orders *Eulypotyphla, Soricomorpha, Erinaceomorpha, Carnivora*, and *Lagomorpha*. Fresh fecal samples were collected and kept individually in tubes containing 500 μL of 1X phosphate-buffered saline (PBS) at -20 °C until they were transported to the laboratory and stored at -80 °C for further processing.

### 2.2. Sample processing and nucleic acids extraction

A previous study showed that many of the samples presented an inhibitory agent that prevented sequencing (Buigues *et al*. 2024). Consequently, only the seventy-six samples lacking such an inhibitory effect were used here (**Supplementary Table S1**). These were combined as previously described into a total of thirteen pools (Buigues *et al*. 2024), each containing between one and twelve samples from the same species (**Supplementary Table S1**). Processing of each pool was carried out as described before (Buigues *et al*. 2024; Carrascosa-Sàez *et al*. 2024). Briefly, an aliquot of each sample was homogenized in a Precellys Evolution tissue homogenizer (Bertin), and the supernatant obtained after centrifugation was filtered using Minisart cellulose acetate syringe filters with a 1.2 µm pore size (Sartorius). For RNA extraction, 250 µL of the filtrate were cleaned with Trizol LS reagent (Invitrogen) and then 280 µL from this step were used for final extraction using the QIAamp Viral RNA minikit (Qiagen).

### 2.3. Sequencing and virus identification

The preparation of libraries from extracted nucleic acids was carried out using the stranded mRNA preparation kit (Illumina), skipping the mRNA enrichment steps and proceeding straight to the fragmentation step. Paired-end sequencing was performed on a NextSeq 550 device with a read length of 150 bp at each end. Raw reads were processed with fastp v0.23.2 (Chen *et al*. 2018), including deduplication, quality filtering, and trimming with a threshold of 20, and those reads below 70 nucleotides in length were removed. De novo sequence assembly was performed using SPAdes v3.15.4 (Antipov *et al*. 2020) with the meta option and MEGAHIT v1.2.9 (Li *et al*. 2015) using default parameters. Then, assembled contigs were clustered to remove replicates using CD-HIT v4.8.1 (Li and Godzik 2006), and those shorter than 1,000 nt were removed. Taxonomic classification was performed using Kaiju v1.9.0 (Menzel, Ng and Krogh 2016) with the subset of the NCBI nr protein database downloaded on June 6, 2023. Viral contigs were identified using Virsorter2 v2.2.4 (Guo *et al*. 2021) and further analyzed with CheckV v1.0.1 (Nayfach *et al*. 2021) for genome quality assessment. Contigs were selected based on size, completeness, and potential vertebrate infectivity.

Coverage statistics for the viral contigs were determined by remapping the filtered and trimmed reads to their corresponding contigs using Bowtie2 v2.2.5 (Langmead and Salzberg 2012). The raw sequence reads were deposited in the Sequence Read Archive (SRA) of GenBank under accession numbers SRR32004501-13. The viral contig described in this study was deposited in GenBank under accession number PQ888429.

### 2.4. Genome annotation and phylogenetic analysis

Protein domains were annotated using Interpro 103.0 (Blum *et al*. 2025). Open reading frames (ORFs) were predicted using ORFfinder (https://www.ncbi.nlm.nih.gov/orffinder). The ORFs corresponding to the viral contig described in this study were compared to NCBI databases using BLASTp to obtain identity values and refine annotations. For phylogenetic analysis at the whole genome level, the forty-eight torovirus sequences currently available in the databases were downloaded. Next, a nucleotide-based multiple alignment was performed using MAFFT v7.505 (Katoh and Standley 2013) and then trimmed with trimAl v1.2rev59 (Capella-Gutiérrez, Silla-Martínez and Gabaldón 2009), setting the gap threshold option at 0.2. The maximum likelihood (ML) phylogenetic trees were obtained with IQ-TREE v2.3.6 (Minh *et al*. 2020), using the built-in ModelFinder function (Kalyaanamoorthy *et al*. 2017) to infer the best substitution model. Branch support was assessed using 1,000 ultra-fast bootstrap replicates (Hoang *et al*. 2018) and 1,000 bootstrap replicates for the SH-like approximate likelihood ratio test.

BEAST v2.7.6 (Drummond and Rambaut 2007) was used to estimate the time of the most recent common ancestor (tMRCA) of toroviruses using the nucleotide sequences of the ORF1b. Nucleotide-based alignment was obtained using MUSCLE v3.8.1551 (Edgar 2004). Sequences showing evidence of recombination by at least three methods implemented in RDP4 (Martin *et al*. 2015) were excluded from the analysis. Additionally, due to a recombination event affecting the entire bovine torovirus clade, the corresponding region (nucleotide positions 6042–6743) was removed from the alignment before further analysis (**Supplementary Table S2)**.

The best-fit nucleotide substitution model was selected using the ModelFinder function implemented in IQ-TREE v2.3.6, which identified GTR+F+I+G4 as the optimal model. Phylogenetic inference was conducted under a strict molecular clock and a coalescent constant-size tree prior. Bayesian Markov Chain Monte Carlo (MCMC) analyses were run for thirty million generations in two independent chains, sampling every 10,000 generations, with a burn-in of 10%. Convergence of parameters was assessed using Tracer v1.7 (Rambaut *et al*. 2018). The maximum clade credibility (MCC) tree was generated after discarding the first 10% of samples using TreeAnnotator v1.10.4 and was visualized with FigTree v1.4.4.

### 2.5. Ethical statement

According to the European directive regulating the protection of animals used for scientific purposes (2010/62/EU, Article 1), subsequently transposed into Spanish legislation (Royal decree 53/213, 1 February, Article 2), procedures used in this study (i.e. capture, non-invasive handling and in situ release of wild animals) are not subject to the condition of animal experimentation and, therefore, an IACUC approval document is not required, but specifically a permit from the competent regional authority.

## 3. Results

From metagenomics analysis, only one viral genome, identified in a fecal sample from a dormouse (*Elyomis quercinus*), was assigned to the order Nidovirales and classified as a torovirus by Kaiju, hereafter designated as Dormouse ToV (DToV). According to the ICTV, the members of the genus *Torovirus* present genomes of about twenty-eight kb in length. Congruently, DToV showed a genome size of 28,555 nt. A total of 8,085 reads from its corresponding library were remapped to this contig with an average coverage of 39.12 ± 13.13. The G+C content was 38.7%, slightly higher than in other genomes, such as bovine, antelope, and porcine toroviruses (38, 37, and 35%, respectively) (Dai *et al*. 2021).

The DToV genome included 5’ and 3’ non-translated regions (NTR, 877 and 199 nt in length, respectively), six characteristic ORFs, and also the two commonly deduced CUG-initiated ORFs encoding U1 and U2 putative proteins within the 5’-NTR and ORF1a, respectively (Ujike and Taguchi 2021) (**Figure 1**; **Supplementary Table S3**). The replicase polyprotein (pp1ab), jointly encoded by ORF1a and ORF1b, included eleven predicted domains: ADP-ribose 1-phosphatase (ADRP), papain-like protease (PLP), 3C-like main protease (M^pro^), cyclic phosphodiesterase (CPD), nidovirus RdRp-associated nucleotidyltransferase (NiRAN), RNA-dependent RNA polymerase (RdRp), Zn-binding domain (ZBD), Helicase, 3’-to 5’ exoribonuclease domain (ExoN), nidoviral uridylate-specific endoribonuclease (NendoU), and ribose 2’-O-methyltransferase (MT). The CPD domain (IPR039573), detected at the 3’-end of ORF1a, is related to NS2, identified only in a lineage A betacoronavirus but at a different genome location (Ujike and Taguchi 2021). In addition, pp1ab also included six peptidase 3C cleavage sites.

**Figure 1.**
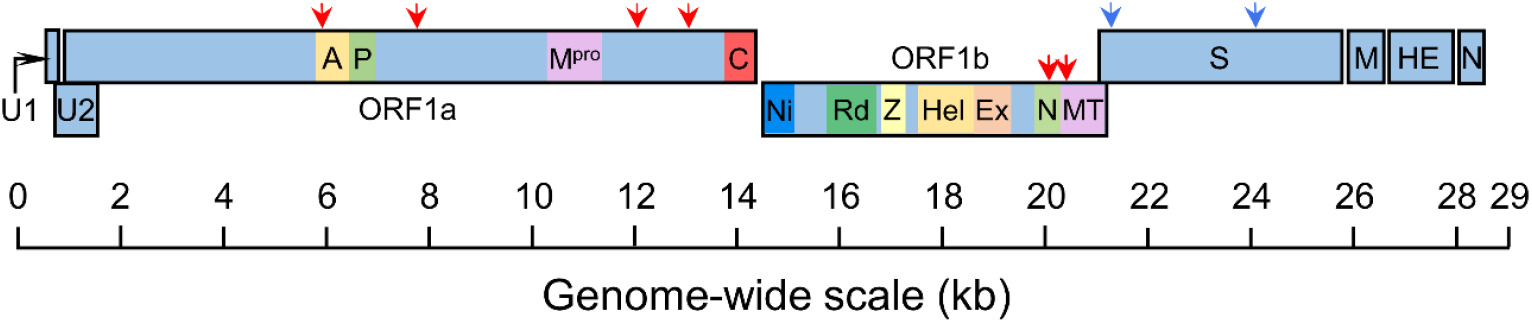
Schematic representation of the DToV genome showing ORF distribution. Conserved protein motifs/domains are highlighted in coloured boxes as follows: A represents the ADP-ribose 1-phosphatase (ADRP); P represents the papain-like protease (PLP); M^pro^ represents the 3C-like main protease; C represents the cyclic phosphodiesterase (CPD); Ni represents the nidovirus RdRp-associated nucleotidyltransferase (NiRAN); Rd represents the RNA-dependent RNA polymerase (RdRp); Z represents the Zn-binding domain (ZBD); Hel represents the Helicase; Ex represents the 3’-to 5’ exoribonuclease domain (ExoN); N represents the nidoviral uridylate-specific endoribonuclease (NendoU); MT represents the ribose 2’-O-methyltransferase. Peptidase 3C and furin cleavage sites are indicated by red and blue arrows, respectively.

The last third of the DToV genome contained genes coding for spike (S), membrane (M), haemagglutinin-esterase (HE), and nucleocapsid (N) proteins (**Figure 1**; **Supplementary Table S3**). ORF1b and S were slightly overlapping, while short intergenic regions were present between the four C-terminal genes. The S protein showed the two typical furin cleavage sites found in toroviruses (Ujike and Taguchi 2021). In addition, this protein lacked the cysteine-rich region between the transmembrane domain and the cytoplasmic tail at the C-terminal end, which is present in coronaviruses but not in toroviruses (Petit *et al*. 2007). Upstream of the M, HE, and N genes, a short intergenic motif (5′-ACN_3-4_CUUUAGA-3′), previously described as a trans-regulatory sequence (TRS), is always detected in toroviruses (Smits *et al*. 2005). This motif showed a small sequence variation upstream of the N gene (5′-ACN_3_-CUUUAGT-3′), which may have implications for the complex transcription mechanism of this group of viruses (Ujike and Taguchi 2021).

To compare genome-wide characteristics between DToV and other toroviruses, a BLAST analysis was performed against different toroviruses, including a representative of the three currently accepted species and other complete or near-complete genomes identified in different hosts. Specifically, sequences from antelope, goat, and Tasmanian devil (Torovirus sp.) toroviruses were used, in addition to one identified in a camel-derived tick (Bangali virus) (Zhang *et al*. 2021), possibly acting as a transmission vector (**Table 1**). Thus, all complete or near-complete torovirus genomes available to date have been identified in ungulates, except for Torovirus sp., detected in a marsupial carnivore (Chong *et al*. 2019). From pairwise comparisons against DToV, nucleotide and amino acid identities were estimated for the six major ORFs. For each ORF, identities were very similar across the different toroviruses, except Bangali virus, which had lower values for ORFs 1b, M and N. Overall, nucleotide and amino acid identities were considerably low, around or below 60% for ORFs 1a, HE and N, below 70% for S, and around 80% for 1b and M. These results showed that DToV was different from the previously described torovirus sequences and that these differences strongly varied across the genome.

**Table 1.** Nucleotide and amino acid (between brackets) identities (%) of six open reading frames between DToV and representative torovirus sequences (accessions are provided between brackets). S, spike glycoprotein; M, membrane protein; HE, haemagglutinin-esterase; N, nucleocapsid protein.

| ORFs | BToV<br>(ON376728) | PToV<br>(MT684462) | EToV<br>(MG996765) | Antelope ToV<br>(MZ438674) | Goat ToV<br>(NC_034976) | Torovirus sp.<br>(MK521914) | Bangali virus<br>(NC_076940) |
| --- | --- | --- | --- | --- | --- | --- | --- |
| 1a | 62.0 (53.5) | 62.3 (53.6) | 62.8 (54.9) | 61.6 (53.9) | 61.9 (54.0) | 62.2 (54.2) | 60.3 (52.6) |
| 1b | 75.8 (80.8) | 76.0 (80.7) | 77.2 (82.4) | 76.0 (79.9) | 76.0 (82.0) | 76.0 (81.8) | 71.0 (74.3) |
| S | 66.9 (68.5) | 65.9 (65.6) | 68.1 (68.2) | 66.3 (66.9) | 66.9 (68.6) | 66.0 (68.6) | 67.0 (67.0) |
| M | 78.1 (86.3) | 75.9 (84.6) | 77.0 (83.8) | 73.6 (84.2) | 78.1 (85.0) | 78.1 (86.3) | 69.4 (68.2) |
| HE | 61.6 (50.7) | 58.0 (50.2) | - | 59.0 (46.4) | 61.6 (51.3) | 60.7 (51.8) | 57.9 (47.4) |
| N | 56.7 (54.9) | 60.6 (56.9) | 62.2 (55.5) | 56.4 (56.7) | 61.1 (55.8) | 61.0 (55.8) | 49.8 (44.3) |

A phylogenetic analysis was performed using all currently available complete torovirus genomes (**Figure 2A**). The tree showed that DToV was clearly distant from the three currently accepted torovirus species and from other unclassified toroviruses identified in different hosts. From the data presented here, DToV emerged as the first complete genome of a rodent-associated torovirus. However, a metagenomic analysis recently identified torovirus sequences in fecal samples from the rodent *Myodes glareolus* species (Raghwani *et al*. 2023). A complete genome could not be assembled in that study, but reanalysis of the raw sequencing data identified eight contigs associated with toroviruses. From these contigs, the complete sequence of the HE and N genes was recovered, and phylogenetic analysis involving these genes showed that DToV remained in a basal position, while the other rodent-associated torovirus was much closer to viruses identified in other hosts (**Figure 2B**). This suggests that virus-host specificity in toroviruses may not be as high as proposed (Flies *et al*. 2023). The main incongruence between the two phylogenetic analyses shown in **Figure 2** corresponds to the position of the antelope ToV, which is explained by its involvement in an HE gene recombination event (Dai *et al*. 2021).

**Figure 2.**
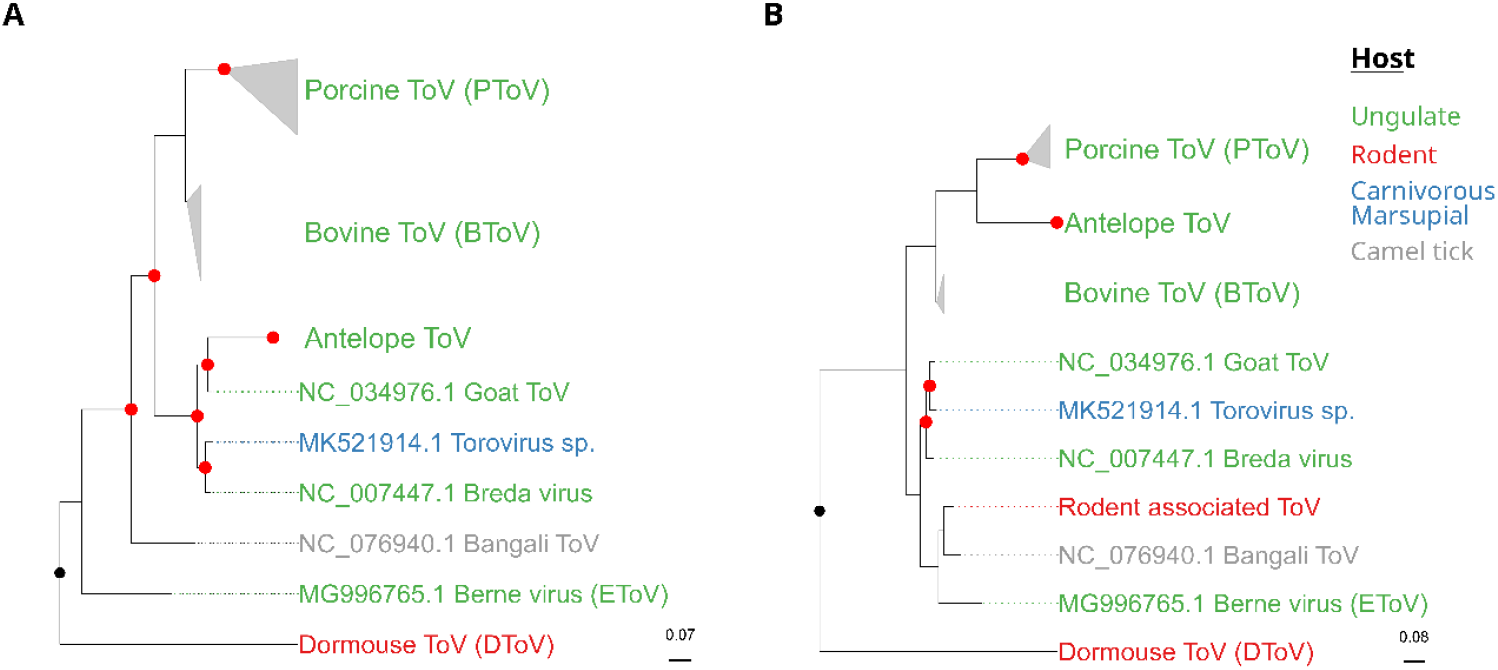
Maximum likelihood phylogenetic trees based on the nucleotide sequence of ToV representatives for the whole genome (A) and the region including HE and N genes (B). The partial sequence corresponding to a recently described rodent-associated ToV (Raghwani *et al*. 2023) was also included in B. Trees were rooted using as outgroup a snake virus sequence (NC_046963) from another genus of the family *Tobaniviridae*. Substitution models used were GTR+F+G4 and TIM2+F+I+G4, respectively. SH-aLRT and ultrafast bootstrap values higher than 80 and/or 95, respectively, are indicated in red circles. The tree is rooted at the midpoint. Scale bars indicate the evolutionary distance in nucleotide substitutions per site.

A Bayesian tree based on complete nucleotide sequences of the ORF1b was obtained to estimate divergence times in ToVs (**Figure 3**). The most recent common ancestor of toroviruses was dated 1,286 years ago (95% HPD=858-1831), a higher value than previously described with the M gene dating analysis (Dai *et al*. 2021), and which reflected the inclusion of the new sequence identified in our study. It is worth mentioning that this analysis was based on ORF1b because it is much longer than the M gene, highly conserved among all toroviruses, and includes the viral polymerase. Nevertheless, the estimated mean substitution rate for this dataset was 3.88·10^-4^ substitutions per site per year (95% HPD = 2.5·10^-4^-5.3·10^-4^), nearly identical to that previously described for the M gene (Dai *et al*. 2021).

**Figure 3.**
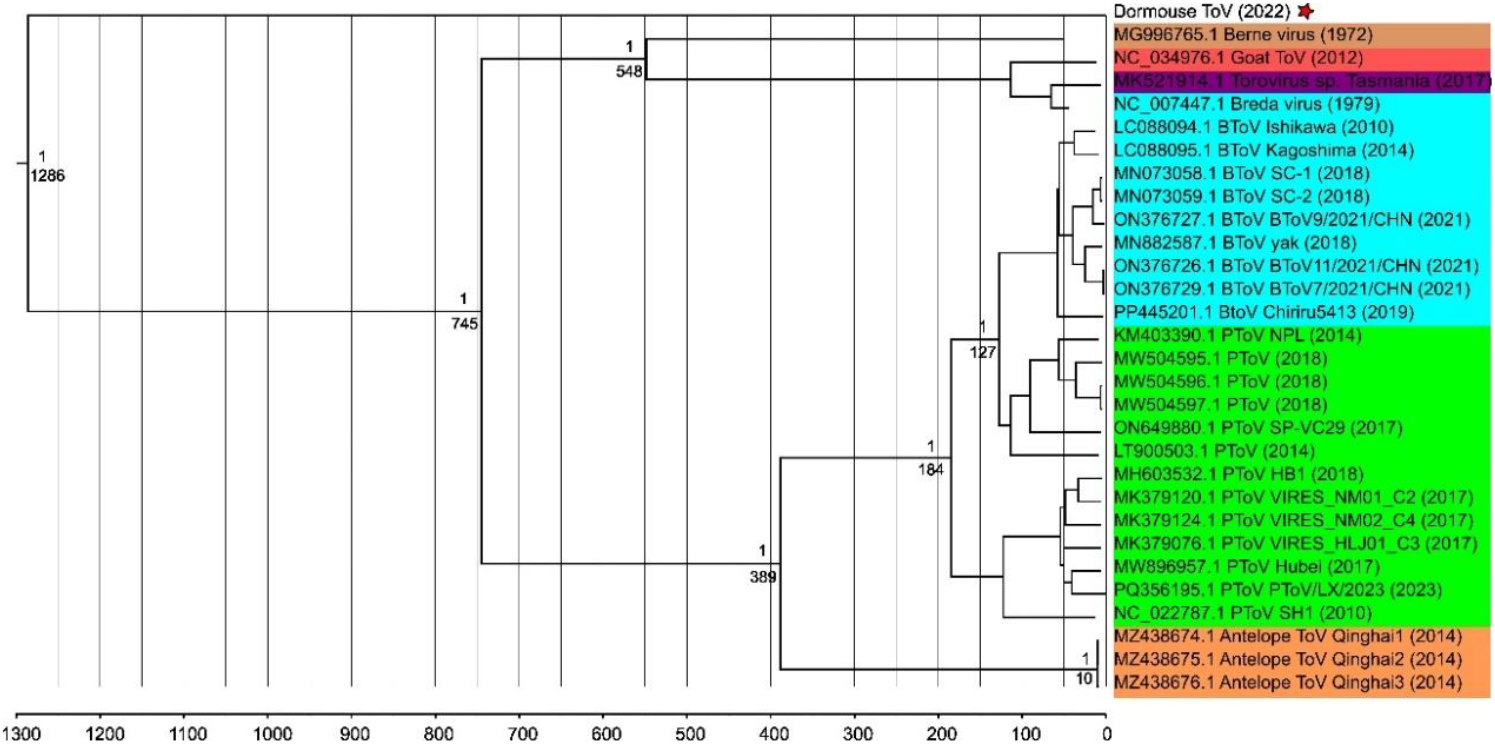
Bayesian phylogenetic tree based on the nucleotide sequence of ToV representatives for the ORF1b. Accession number, species and strain name, and year of sampling are on the tip labels. Node labels indicate the posterior probabilities and the time of the most recent common ancestor (tMRCA). DToV was marked with a red star.

## 4. Discussion

The diversity of toroviruses is probably much greater than currently considered by the ICTV. Only three species are defined in ungulate hosts (BToV, EToV, and PToV), although toroviruses appear to infect a wide range of animal species. This is supported by indirect evidence, such as the detection of antibodies against toroviruses in the sera of other animals, including goats, sheep, rabbits, and mice (Weiss *et al*. 1984; Brown, Beards and Flewett 1987). Moreover, electron microscopy studies have detected torovirus-like particles in cat and human faeces (Beards *et al*. 1984; Muir *et al*. 1990). However, direct characterisation requires the identification of viral sequences, ideally complete genomes. The majority of torovirus complete genome sequences are of porcine and bovine origin, potentially as a result of sampling bias. As new sequences are added to the databases, it is noted that toroviruses are not exclusive to ungulates, since they have already been observed in other taxonomic groups, such as in a marsupial carnivore (Chong *et al*. 2019) and the one detected in this work in rodents. This will probably lead to the definition of new species and probably also new subgenera. For instance, Bangali virus has been recently included into a new subgenus (*Bantovirus*) within the genus *Torovirus* according to the Virus Taxonomy 2023 Release, while the other toroviruses have been assigned to the subgenus *Renitovirus*. In this context, therefore, it is likely that DToV should be considered not only a new species, but at least a new subgenus.

Although coronaviruses and toroviruses are relatively close groups (Ujike and Taguchi 2021), the latter have been the subject of fewer studies, probably because they are not currently associated with human diseases. Nevertheless, BToVs cause economic losses due to calf diarrhoea (Woode *et al*. 1982; Duckmanton *et al*. 1998), while PToVs seem to cause less severe symptoms, although these may be worsened by co-infections (Ujike and Taguchi 2021). In any case, new evidence supports the zoonotic potential of toroviruses. First, the detection of a torovirus in an insect suggests the possibility of vector-borne transmission (Flies *et al*. 2023). Second, numerous cases of recombination have been observed, not only between different toroviruses but also with viruses of other genera, which can either enhance their pathogenicity or facilitate unexpected host adaptation (Smits *et al*. 2003; Ito *et al*. 2016; Conceição-Neto *et al*. 2017; Lee and Lee 2019). Third, wild ungulates have been identified as potential hosts of undiscovered zoonotic viruses (Olival *et al*. 2017). Indeed, the detection of an antelope torovirus provides evidence of cross-species transmission between wild and domestic ungulates (Dai *et al*. 2021). Finally, the identification of new toroviruses in wild animals, especially rodents, further underscores their cross-species transmissibility and suggests some degree of zoonotic potential, as the proximity of rodents to humans and farm animals may facilitate host-jumping events (Flies *et al*. 2023). In this regard, the remarkable divergence time between DToV and the rest of the sequences makes it tempting to speculate that rodents may be the ancestral host of toroviruses.

Apart from their apparently low pathogenicity, little is known about toroviruses because they have not been cultured, with a few exceptions. In the mid-1980s, EtoV (Berne virus) was the first torovirus to be propagated in cell cultures (Weiss and Horzinek 1986). Since then, no other EToVs have been isolated, suggesting that Berne virus may be a mutant adapted to growth in cultures (Koopmans and Horzinek 1994). More recently, several BToVs have been propagated in cell cultures (Kuwabara *et al*. 2007; Aita *et al*. 2012; Ito *et al*. 2016), which showed, for example, that HE protein is not essential for viral growth under these conditions. BToV is also the only torovirus for which a reverse genetics system is available (Ujike *et al*. 2022), which is likely to become an essential tool for studying important viral aspects such as pathogenesis and vaccine development. In this scenario, studying the S protein, which determines cellular and tissue tropism and host range (Tortorici and Veesler 2019), is crucial to characterise the zoonotic potential of toroviruses. The use of viral pseudotypes has recently allowed us to characterise the spike protein of dozens of viruses from multiple RNA virus families (Dufloo *et al*. 2025). Unfortunately, our attempt to obtain a VSV pseudotype to study the cell tropism of a torovirus was unsuccessful. In the future, other approaches, such as those based on obtaining recombinant viruses (Marqués *et al*. 2024), may help to shed light on the zoonotic potential of toroviruses.

## Conclusions

To date, most of the known torovirus species have been isolated from ungulates (Dai *et al*. 2021). However, the complete genome reported here was recovered from a faecal sample of a Spanish dormouse, suggesting that the host range of toroviruses is broader than currently described. The basal phylogenetic position of the newly described torovirus led us to speculate that rodents may serve as a reservoir for this viral genus. Although the zoonotic potential of toroviruses may be low, some of their characteristics, such as their tendency to recombine and possible vector-borne transmission, point to the need to include them in surveillance programs. Further efforts to understand the infectivity determinants of this group of viruses will be necessary to better characterize their potential for host-jumping events.

## Data availability

Data available in supplementary material.

## Supplementary material

Supplementary Table S1. Small mammal species, collection sites, pooling, and inhibitory. potential of each sample

Supplementary Table S2. Recombination events detected in the ORF1b alignment.

Supplementary Table S3. Genome annotation of DToV.

